## Supplementary Material for "Shaping the physical world to our ends: The left PF technical-cognition area"

**This PDF file includes:**

Fig. S1 and S2

Tables S1 to S7

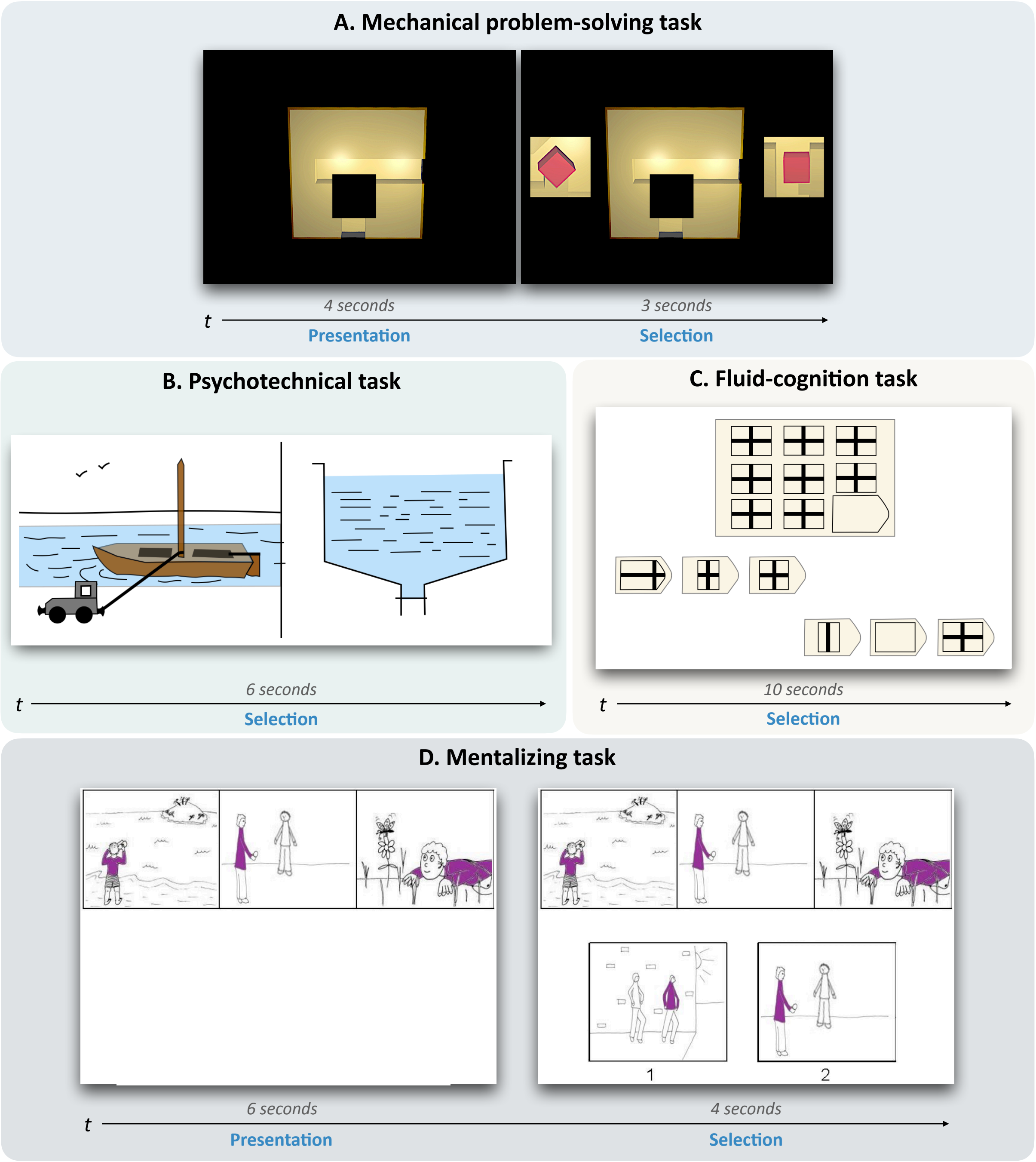


**Fig. S1. Control conditions of the experimental tasks. (A)** In the mechanical problem-solving task (Experiment 1), participants scrutinized the 3D glass box for 4 seconds and then had 3 seconds to decide which of the two missing pieces presented was the correct one to fill the mask. **(B)** In the psychotechnical task (Experiment 2), two situations were displayed for 6 seconds. Participants had to select which of the two displayed situations contained a white square. **(C)** In the fluid-cognition task (Experiment 2), the participants had to select the line of options with the correct one. Contrary to the experimental condition, the control condition only required visual completion. **(D)** In the mentalizing task (Experiment 2), the superior part of the board was shown for 6 seconds. Then the bottom part was presented for 4 additional seconds, with the top part remaining on display. The participants had to select which cartoon was already present in the first three ones. Birgit Völlm gave us the permission to reproduce the pictures in **(D)**. No permission was needed for the pictures in **(A)** as we built this task. The items of the psychotechnical task **(B)** and the fluid-cognition task **(C)** are adapted from commercialized tests and do not correspond to the original items of these tests. For more information, see the Methods section.

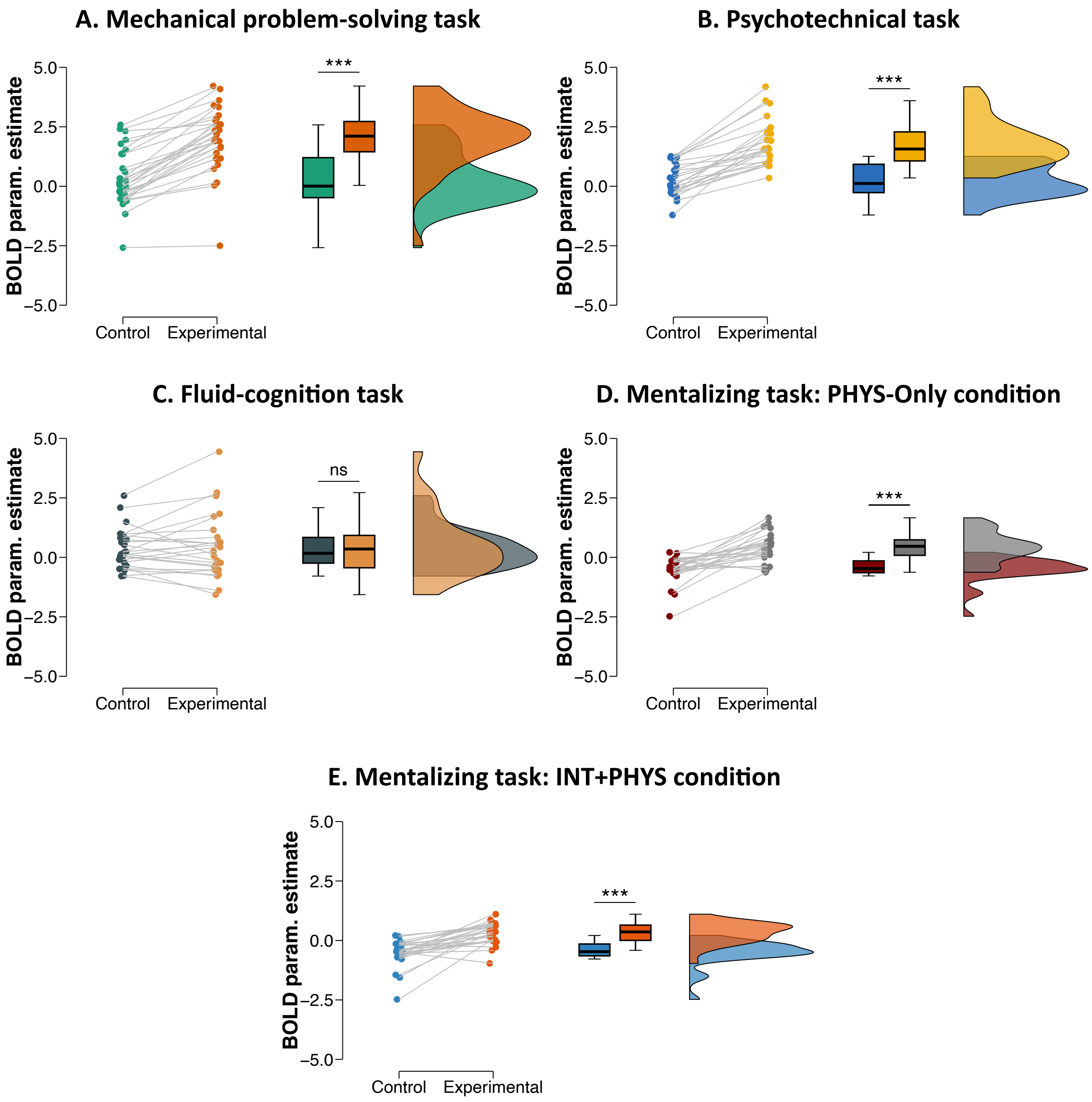


**Fig. S2. ROI univariate results (left area PF) for participants who performed at or below 50%.** BOLD param. estimate refers to the mean BOLD parameter estimate. Boxplots indicate the upper quartile, median and lower quartile, with whiskers extending to the most extreme data point that is no more than 1.5 times the interquartile range from the edge of the box. Bars represent the means and 95% confidence intervals. ns, not significant; ***, $p < .001$.

**Table S1. Local maxima of activation clusters (MNI stereotactic coordinates) for the Mechanical problem-solving task (Experimental condition > Control condition)**

| Cluster size | Hemisphere | Brain region | Peak coordinates | | | *t*-value |
|---|---|---|---|---|---|---|
| | | | *x* | *y* | *z* | |
| 1963 | Left | Supramarginal gyrus (PF) | -59 | -30 | 41 | 11.69 |
| | | Supramarginal gyrus (PF) | -59 | -32 | 31 | 11.31 |
| | | Superior parietal cortex | -8 | -57 | 66 | 11.05 |
| 696 | Left | Dorsal premotor cortex | -20 | -9 | 54 | 14.98 |
| | | Dorsal premotor cortex | -18 | 7 | 66 | 10.14 |
| | | Dorsal premotor cortex | -18 | -7 | 71 | 9.52 |
| 246 | Left | Inferior frontal gyrus (triangular part) | -47 | 37 | 6 | 9.05 |
| | | Anterior prefrontal cortex | -45 | 41 | 20 | 8.86 |
| | | Dorsolateral prefrontal cortex | -41 | 37 | 13 | 7.89 |
| 186 | Left | Inferior frontal gyrus (opercular part) | -50 | 7 | 20 | 9.89 |
| 303 | Right | Cerebellum | 26 | -57 | -54 | 9.71 |
| | | Cerebellum | 28 | -50 | -26 | 8.96 |
| | | Cerebellum | 40 | -48 | -35 | 8.64 |

These results are also illustrated in Fig. 2A. PF, parietal area F.

**Table S2. Local maxima of activation clusters (MNI stereotactic coordinates) for the Psychotechnical task (Experimental condition > Control condition)**

| Cluster size | Hemisphere | Brain region | Peak coordinates | | | *t*-value |
|---|---|---|---|---|---|---|
| | | | *x* | *y* | *z* | |
| 1196 | Left | Supramarginal gyrus (PF) | -57 | -32 | 38 | 13.94 |
| | | Superior parietal cortex | -20 | -69 | 48 | 10.49 |
| | | Intraparietal sulcus | -38 | -34 | 41 | 10.31 |
| 783 | Left | Lateral occipitotemporal cortex | -50 | -66 | -1 | 14.54 |
| | | Lateral occipitotemporal cortex | -41 | -66 | -1 | 11.28 |
| | | Lateral occipitotemporal cortex | -43 | -44 | -15 | 8.16 |
| 216 | Left | Dorsal premotor Cortex | -22 | -7 | 54 | 11.59 |
| 213 | Left | Inferior frontal gyrus (opercular part) | -50 | 7 | 27 | 8.50 |
| 2629 | Right | Superior parietal cortex | 24 | -62 | 50 | 12.4 |
| | | Lateral occipitotemporal cortex | 44 | -62 | -8 | 11.62 |
| | | Intraparietal sulcus | 37 | -39 | 45 | 11.23 |
| 298 | Right | Inferior frontal gyrus (opercular part) | 49 | 9 | 27 | 9.77 |
| | | Inferior frontal gyrus (opercular part) | 49 | 9 | 18 | 8.50 |
| 230 | Right | Dorsal premotor cortex | 24 | 0 | 52 | 8.65 |
| | | Dorsal premotor cortex | 30 | 2 | 66 | 7.49 |

These results are also illustrated in Fig. 2B. PF, parietal area F.

**Table S3. Local maxima of activation clusters (MNI stereotactic coordinates) for the Fluid-cognition task (Experimental condition > Control condition)**

| Cluster size | Hemisphere | Brain region | Peak coordinates | | | *t*-value |
|---|---|---|---|---|---|---|
| | | | *x* | *y* | *z* | |
| 1147 | Left | Dorsal premotor cortex | -29 | 0 | 59 | 11.64 |
| | | Dorsolateral prefrontal cortex | -47 | 25 | 29 | 10.67 |
| | | Inferior frontal gyrus (opercular part) | -45 | 7 | 31 | 10.08 |
| 127 | Left | Thalamus | -13 | -25 | 13 | 8.99 |
| | | Thalamus | -6 | -12 | 6 | 7.86 |
| | | Thalamus | -20 | -30 | 4 | 7.85 |
| 140 | Left | Insula | -31 | 23 | 2 | 12.36 |
| 974 | Right | Inferior frontal gyrus (opercular part) | 40 | 11 | 31 | 11.95 |
| | | Inferior frontal gyrus (opercular part) | 49 | 21 | 31 | 10.90 |
| | | Inferior frontal gyrus | 46 | 7 | 31 | 10.15 |
| 661 | Right | Dorsal premotor cortex | 30 | 2 | 57 | 13.25 |
| | | Dorsal prefrontal cortex | 30 | 11 | 50 | 9.73 |
| | | Dorsal prefrontal cortex | 24 | 21 | 50 | 9.04 |
| 201 | Right | Insula | 33 | 21 | 4 | 13.63 |
| 9887 | Left/Right | Cerebellum | -8 | -73 | -24 | 16.08 |
| | | Cerebellum | 3 | -71 | -24 | 15.84 |
| | | Occipital cortex | -38 | -76 | -12 | 14.88 |
| 806 | Left/Right | Medial superior frontal cortex | -6 | 11 | 52 | 11.49 |
| | | Medial superior frontal cortex | 8 | 16 | 45 | 10.77 |
| | | Medial superior frontal cortex | -6 | 27 | 43 | 9.82 |
| 644 | Left/Right | Thalamus | 5 | -25 | -5 | 13.45 |
| | | Thalamus | -4 | -28 | -5 | 12.63 |
| | | Thalamus | 10 | -23 | 13 | 9.94 |

These results are also illustrated in Fig. 2C. Fig. 2C shows activation in the superior parietal cortices (SPC) and lateral occipitotemporal cortices (LOTC), which could seem not to be reported here. This activation belongs to the cluster with the size of 9887, which is a very large cluster.

**Table S4. Local maxima of activation clusters (MNI stereotactic coordinates) for the Mentalizing task (PHYS-Only condition > Control condition)**

| Cluster size | Hemisphere | Brain region | Peak coordinates | | | *t*-value |
|---|---|---|---|---|---|---|
| | | | *x* | *y* | *z* | |
| 274 | Left | Lateral occipitotemporal cortex | -50 | -62 | 4 | 10.27 |
| | | Lateral occipitotemporal cortex | -54 | -66 | -5 | 8.91 |
| | | Lateral occipitotemporal cortex | -45 | -69 | -1 | 8.75 |
| 133 | Left | Supramarginal gyrus (PF) | -59 | -25 | 34 | 8.44 |
| | | Supramarginal gyrus (PF) | -57 | -34 | 41 | 7.97 |
| | | Supramarginal gyrus (PF) | -52 | -37 | 34 | 7.90 |
| 267 | Right | Lateral occipitotemporal cortex | 46 | -60 | -3 | 9.97 |
| | | Lateral occipitotemporal cortex | 51 | -53 | -1 | 8.22 |
| | | Lateral occipitotemporal cortex | 49 | -64 | 6 | 6.63 |
| 138 | Right | Supramarginal gyrus | 60 | -28 | 48 | 7.96 |
| | | Supramarginal gyrus | 58 | -23 | 34 | 7.62 |
| | | Supramarginal gyrus | 51 | -32 | 50 | 6.70 |

These results are also illustrated in Fig. 2D. PF, parietal area F.

**Table S5. Local maxima of activation clusters (MNI stereotactic coordinates) for the Mentalizing task (INT+PHYS condition > Control condition)**

| Cluster size | Hemisphere | Brain region | Peak coordinates | | | *t*-value |
|---|---|---|---|---|---|---|
| | | | *x* | *y* | *z* | |
| 1464 | Left | Lateral occipitotemporal cortex | -50 | -71 | -3 | 13.28 |
| | | Lateral occipitotemporal cortex | -43 | -46 | -21 | 10.72 |
| | | Angular gyrus | -54 | -64 | 8 | 10.59 |
| 256 | Left | Cerebellum | -18 | -71 | -28 | 10.48 |
| | | Cerebellum | -15 | -78 | -42 | 9.57 |
| 122 | Left | Supramarginal gyrus (PF) | -59 | -32 | 31 | 9.02 |
| 1752 | Right | Lateral occipitotemporal cortex | 46 | -60 | 2 | 14.35 |
| | | Lateral occipitotemporal cortex | 51 | -64 | -5 | 11.72 |
| | | Angular gyrus | 42 | -55 | 15 | 11.13 |
| 282 | Right | Temporal pole | 51 | 2 | -24 | 9.60 |
| | | Middle temporal gyrus | 51 | -5 | -19 | 9.13 |
| | | Temporal pole | 49 | 18 | -28 | 8.74 |
| 165 | Right | Inferior frontal gyrus (triangular part) | 51 | 30 | 8 | 8.42 |
| | | Inferior frontal gyrus (opercular part) | 44 | 7 | 20 | 7.28 |
| | | Inferior frontal gyrus (opercular part) | 51 | 18 | 25 | 6.23 |

These results are also illustrated in Fig. 2E. PF, parietal area F.

**Table S6. Local maxima of activation clusters (MNI stereotactic coordinates) for the Mentalizing task (INT+PHYS condition > PHYS-Only condition)**

| Cluster size | Hemisphere | Brain region | Peak coordinates | | | *t*-value |
|---|---|---|---|---|---|---|
| | | | *x* | *y* | *z* | |
| 235 | Left | Angular gyrus | -50 | -62 | 25 | 8.56 |
| | | Angular gyrus | -43 | -57 | 27 | 7.83 |
| | | Angular gyrus | -57 | -50 | 13 | 7.66 |
| 310 | Right | Angular gyrus | 49 | -50 | 18 | 9.15 |
| | | Lateral occipitotemporal cortex | 51 | -60 | 15 | 8.99 |
| | | Lateral occipitotemporal cortex | 42 | -50 | 13 | 7.71 |
| 212 | Right | Middle temporal gyrus | 53 | -2 | -17 | 8.45 |
| | | Temporal pole | 56 | 14 | -26 | 7.39 |
| | | Temporal pole | 51 | 7 | -28 | 7.32 |
| 163 | Left/Right | Medial prefrontal cortex | -4 | 53 | 29 | 8.22 |
| | | Medial prefrontal cortex | 3 | 53 | 20 | 7.76 |

These results are also illustrated in Fig. 2F.

**Table S7. Local maxima of activation clusters (MNI stereotactic coordinates) for the conjunction analysis (Mechanical problem-solving AND Psychotechnical AND INT+PHYS AND PHYS-Only)**

| Cluster size | Hemisphere | Brain region | Peak coordinates | | |
|---|---|---|---|---|---|
| | | | *x* | *y* | *z* |
| 374 | Left | Supramarginal gyrus (PF) | -56 | -29 | 36 |

These results are also illustrated in Fig. 2G. PF, parietal area F.
